# Modeling Attention Control Using A Convolutional Neural Network Designed After The Ventral Visual Pathway

**DOI:** 10.1101/473124

**Authors:** Chen-Ping Yu, Huidong Liu, Dimitris Samaras, Gregory Zelinsky

**Affiliations:** Phiar Technologies, Inc., Palo Alto, CA; Stony Brook University, Stony Brook, NY

## Abstract

Recently we proposed that people represent object categories using category-consistent features (CCFs), those features that occur both frequently and consistently across a categorys exemplars [70]. Here we designed a Convolutional Neural Network (CNN) after the primate ventral stream (VsNet) and used it to extract CCFs from 68 categories of objects spanning a three-level category hierarchy. We evaluated VsNet against people searching for the same targets from the same 68 categories. Not only did VsNet replicate our previous report of stronger attention guidance to subordinate-level targets, with its more powerful CNN-CCFs it was able to predict attention control to individual target categories–the more CNN-CCFs extracted for a category, the faster gaze was directed to the target. We also probed VsNet to determine where in its network of layers these attention control signals originate. We found that CCFs extracted from VsNet’s V1 layer contributed most to guiding attention to targets cued at the subordinate (e.g., police car) and basic (e.g., car) levels, but that guidance to superordinate-cued (e.g., vehicle) targets was strongest using CCFs from the CIT+AIT layer. We also identified the image patches eliciting the strongest filter responses from areas V4 and higher and found that they depicted representative parts of an object category (e.g., advertisements appearing on top of taxi cabs). Finally, we found that VsNet better predicted attention control than comparable CNN models, despite having fewer convolutional filters. This work shows that a brain-inspired CNN can predict goal-directed attention control by extracting and using category-consistent features.

## 1. Introduction

### Overview

The brain’s ability to flexibly exert a top-down control over motor behavior is fundamentally important for the achievment of visuomotor goals and the performance of everyday tasks (Hayhoe and Ballard refs). It does this with extreme efficiency, and at times seemingly without effort. Yet, despite being a core cognitive process affecting much of our behavior, a neurocomputational understanding of goal-directed attention control is still in its infancy. Existing computational models of attention control are either relatively narrow in scope or restricted in the types of inputs that they can accept [13, 22, 43]. Here we introduce VsNet, a neurocomputational model of attention control designed after the primate ventral stream of visually-responsive brain areas.

VsNet advances existing models of attention control in several respects. First, and most practically, it is image computable, meaning that it can accept the same visually complex and unlabelled imagery that floods continuously into the primate visual system. This is essential for a model aimed at understanding attention control in the real world, as objects in our perceptual experience do not come with labels telling us what and where they are. Second, VsNet is the first convolutional neural network (CNN) model of attention control. CNNs, one class of artifical deep neural network, have been setting new benchmarks over diverse domains, not the least of which is the automated (without human input) recognition of visually-complex categories of objects [30, 50, 56, 17]. A third and core source of VsNet’s capacity to predict attention control is its extraction of visual features from image examplars that are most represenetative of an object category, a topic that we discuss in detail below. In short, VsNet harnesses the power of deep learning to extract the category-consistent features used by the network of brain areas controlling the goal-directed application of attention.

However, what distinguishes VsNet the most is that it is a brain-inspired CNN of attention control. Our approach is neurocomputational in that, given the many ways that models of attention control could be built, we look to the rich neuroscience literature for design inspiration and parameter specification. Most broadly, VsNet is a multi-layered deep network, making its architecture analogous to the architecture of brain structures existing along the ventral pathway. The brain’s retinotopic application of filters throughout most of these ventral areas also embody a parallelized convolution similar to unit activation across the layers of a CNN [26, 65, 4, 18]. This parallel between a CNN and the organization of the ventral stream has not gone unnoticed [29], and unit activation across the layers of a CNN has even been used to predict neural activity recorded from brain areas in response to the same image content [4, 66]. VsNet extends this work by making the architecture of its levels also brain-inspired, each modeled after a specific brain area in the primate ventral stream. In contrast, existing neurocomputational efforts have used either AlexNet [30] or one of its feed-forward variants [72, 56, 58], which are pre-trained CNNs designed purely to win image classification competitions (e.g., the ILSVRC2012 challenge, also known as the ImageNet dataset, [50]) without regard for the structural and functional organization of the primate ventral visual system. The same disregard for neurobiological constraint applies to later generations of deep networks using different architectures [17, 71, 19]. Determining how VsNet’s performance compares to less brain-inspired CNNs is one broad aim of our study. Another broad aim is to predict the goal-directed allocation of overt attention as people search for categories of objects, as discussed next.

### Categorical Search and Category-Consistent Features

CNNs have been used to predict the bottom-up allocation of attention to scenes [20, 32, 64], but they have not been used to model the top-down control of attention. VsNet is the first. We demonstrate this by predicting the degree that eye movements made by human participants are guided to targets in a categorical search task. The spatial locations fixated via eye movements make the ideal behavioral ground truth for our purpose, as an eye movement is the most basic observable behavior widely believed to indicate a shift of spatial attention [55, 40]. Categorical search is the search for a target that is designated only by its object category. This task can be contrasted with the more common exemplar search task, where participants are cued with an image showing the exact object that they are to search for. Categorical search is therefore perfect for studying the goal directed control of attention in a quasi-realistic context, one where perfect knowledge of a target’s appearance is not assumed. While a historically neglected task (see [74], for discussion), recent research has revealed several important properties of categorical search. Most fundamentally, attention *can* be guided to target categories, as exemplified by the above-chance direction of initial search saccades to target category exemplars in search arrays [67]. Subsequent work has shown that: the strength of the control signal guiding attention to categorical targets depends on the amount of target-defining information provided in the category cue (e.g., stronger guidance for work boot than footwear) [37], that search is guided to distractors that are visually similar to the target category (guidance to a hand fan when searching for a butterfly) [75], that guidance improves with target typicality (stronger guidance to an office chair than a lawn chair) [36], and that guidance becomes weaker as targets climb the category hierarchy (the guidance to “race car” is greater than the guidance to “car”, which is greater than that to “vehicle”) [70]. It is this latter effect of category hierarchy on attention control that is the manipulation of interest in the present study.

In previous work we used a generative model to predict the strength of categorical search guidance across the subordinate (e.g., taxi), basic (e.g., car), and superordinate (e.g., vehicle) levels of a category hierarchy [70]. Specifically, SIFT [35] and color histogram features were extracted from 100 image exemplars of 48 object categories, and the Bag-of-Words (BoW, [6]) method was used to put these features into a common feature space. We then selected those features that are visually most representative of each of these categories, what we termed to be their *Category-Consistent Features* (CCFs). Specifically, for each BOW feature we obtained its responses to all the images of each of a category’s exemplars, averaged these responses over the exemplars, and then divided this average by the standard deviation in responses to obtain a feature-specific Signal-to-Noise Ratio (SNR). A feature having a high SNR would therefore be one that occurred both frequently and consistently across a category’s exemplars. Clustering the features’ SNRs and selecting only the highest, we obtained the CCFs for each of the target categories. Using this BoW-CCF model we were able to predict how behavioral performance was affected by target specification at the three levels of the category hierarchy. One effect was what we called the “subordinate-level advantage” in target guidance; the time that it took gaze to first land on the target (time-to-target) increased with movement up the hierarchy. This result is consistent with related work showing behavioral benefits linked to more detailed and precise visual working memory templates [52]. We showed that a simple count of the number of CCFs for object categories at each level captured almost perfectly the subordinate-level advantage observed in categorical search; more CCFs were selected for categories at the subordinate level than either the basic or superordinate levels. We interpreted this result as indicating that the greater number of CCFs used to represent subordinate-level targets resulted in more detailed visual working memory templates for these target categories that can be used in attention control. The original source should be consulted for more details [70].

The BoW-CCF model offered the first computationally explicit explanation for how level in a category hierarchy affects attention control, but its predictive power in [70] was limited to the three hierarchical levels, essentially three data points. Moreover, in recent years deep learning methods have largely made the once popular BoW method obsolete, due to their significantly better performance in large scale image classification [51, 30], as well as their rich feature representations [47]. The present study extends our previous work in an important respect; rather than predicting the overall effect of hierarchical level on attention control, we now attempt to predict the level of attention control for individual target categories across this same three-level hierarchy. We tried and failed to do this using the BoW-CCF model, but by using VsNet to extract CCFs we show that this audacious goal is in fact attainable.

## 2. General Methods

### Extracting CNN-CCFs

The CCF method selects representative features (may or may not be discriminative) that appear both frequently and consistently across the exemplars of an object category, but the method itself is largely feature independent. In our previous work we selected CCFs from a large pool of BoW features; in our current adaptation we select CCFs from an even larger pool of features from a trained CNN (see Materials and Methods for details regarding model training), where each trained filter is considered a feature and a potential CCF. We hypothesize that the more powerful CNN-CCF features will represent more meaningful visual dimensions of an object category; whereas BoW-CCFs might have coded the fact that many taxis are yellow and represented the various intensity gradiants associated with their shape, a CNN-CCF representation of taxis might additionally capture tires, headlights, and the signs typically mounted to their roofs. We further hypothesize that these richer feature representations, to the extent that they are psychologically meaningful, will allow for better predictions of attention behavior.

The specific CNN-CCF selection process is illustrated in Figure 1 for the taxi category and a hypothetical network. Given an object category with *n* exemplars of size *m* × *m*, and a trained CNN with *L* convolutional layers each containing *K* filters, we forward pass all exemplars through the network to obtain an activation profile of size *m* × *m* × *n* for every convolutional filter, 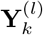, where *l* and *k* are indices to the layer and filter number, respectively. Each 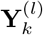 is then reduced to a 1 × *n* vector, 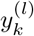, by performing global sum-pooling over each image’s *m*×*m* activation map. This pooling yields the overall activation of each filter in response to an exemplar image. Having these exemplar-specific filter responses, we then borrow from the BoW-CCF pipeline and compute an SNR for each filter:

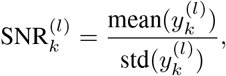

where the mean and standard deviation are computed over the exemplars. Applying this equation to the activation profile from each filter produces a distribution of SNRs. Higher SNRs would indicate stronger and more consistent filter responses, making these filters good candidates for being CCFs. To identify these CCFs we fit a two-component Gamma-Mixture-Model to the SNR distribution, a method similar to Parametric Graph Partitioning [68, 69]. We use a Gamma distribution because it has been shown to model spiking neuron activity [34, 33], and we have observed that it describes our CNN SNR distributions very well. The CCFs are then defined as the filters having SNRs higher than the crossover point of the two Gamma components. This pipeline for extracting CNN-CCFs was applied on each convolutional layer independently, as filter activations have different ranges at different layers.

**Figure 1.**
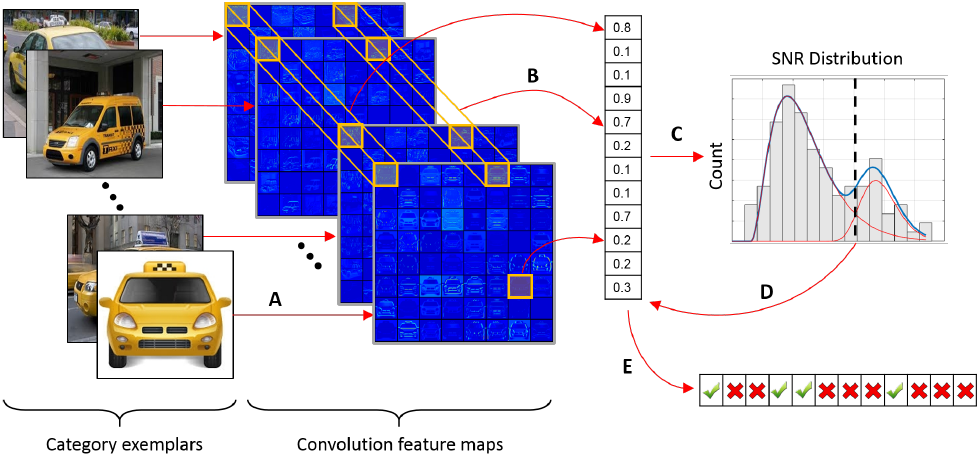
Pipeline of the CNN-CCF extraction method. A. A set of category exemplars, in this case images of taxis, are input into a trained CNN. B. Activation maps (or feature maps) in response to each exemplar are obtained for every convolutional filter at each layer. Shown are 64-cell activation maps in a hypothetical layer, where each cell indicates a convolutional filter’s response to a given exemplar. In this example, 64 SNRs would be computed (12 shown) by analyzing activation map values for each of the 64 filters across the taxi exemplars. C. A two-component Gamma mixture model is fit to the distribution of SNRs, and the cross-over point determines the CCF selection threshold. E. Filters having SNRs above this threshold are retained as the CCFs for a given category; filters having below-threshold SNRs are pruned away.

### Designing and Comparing Brain-Inspired CNNs

There are many good reasons why deep neural networks should be designed more closely after the primate brain. For example, network design has focused on architectures meant to improve network performance, but these are largely ad hoc and not strongly driven by theory. Our broad perspective is that the brain may already have found the best model design, at least for the basic task of visual object classification, and that we need only to consult the voluminous work on brain organization to learn how to implement this design as a deep network.

VsNet is a rough first attempt to build a brain-inspired deep neural network. This effort is “rough” because the neural constraints that we introduce relate only to the gross organization of brain areas along the primate ventral visual stream. There are far more detailed levels of system organization that might also be considered, but as a first pass we decided to focus on only the gross network architecture. We believe that this level would likely reveal the greatest benefit of a brain-inspired design, with the belief that future, more detailed brain-inspired models will only get better. Specifically, we designed VsNet to reflect four widely accepted and highly studied properties of the ventral pathway. First, VsNet’s five convolutional layers are mapped to the five major ventral brain structures [28, 8, 39, 27, 54]. VsNet has a V1, a V2, a combined hV4 and LOC1/2 layer that we refer to as V4-like, a PIT, and a CIT/AIT layer, with these five convolutional layers followed by two fully-connected classification layers. Second, the number of filters in each of VsNet’s five convolutional layers are proportional to the number of neurons, estimated by brain surface area [44, 10], in the corresponding five brain structures. Third, the range of filter sizes at each level layer are informed by the range of receptive field sizes for visually responsive neurons in the corresponding structures. And fourth, VsNet differs from other strictly feedforward architectures in that it adopts a brain-inspired implementation of bybass connections based on what is known about the connectivity between layers in the primate ventral visual stream. Figure 2 and Materials and Methods (MM) should be consulted for additional architectural design details.

**Figure 2.**
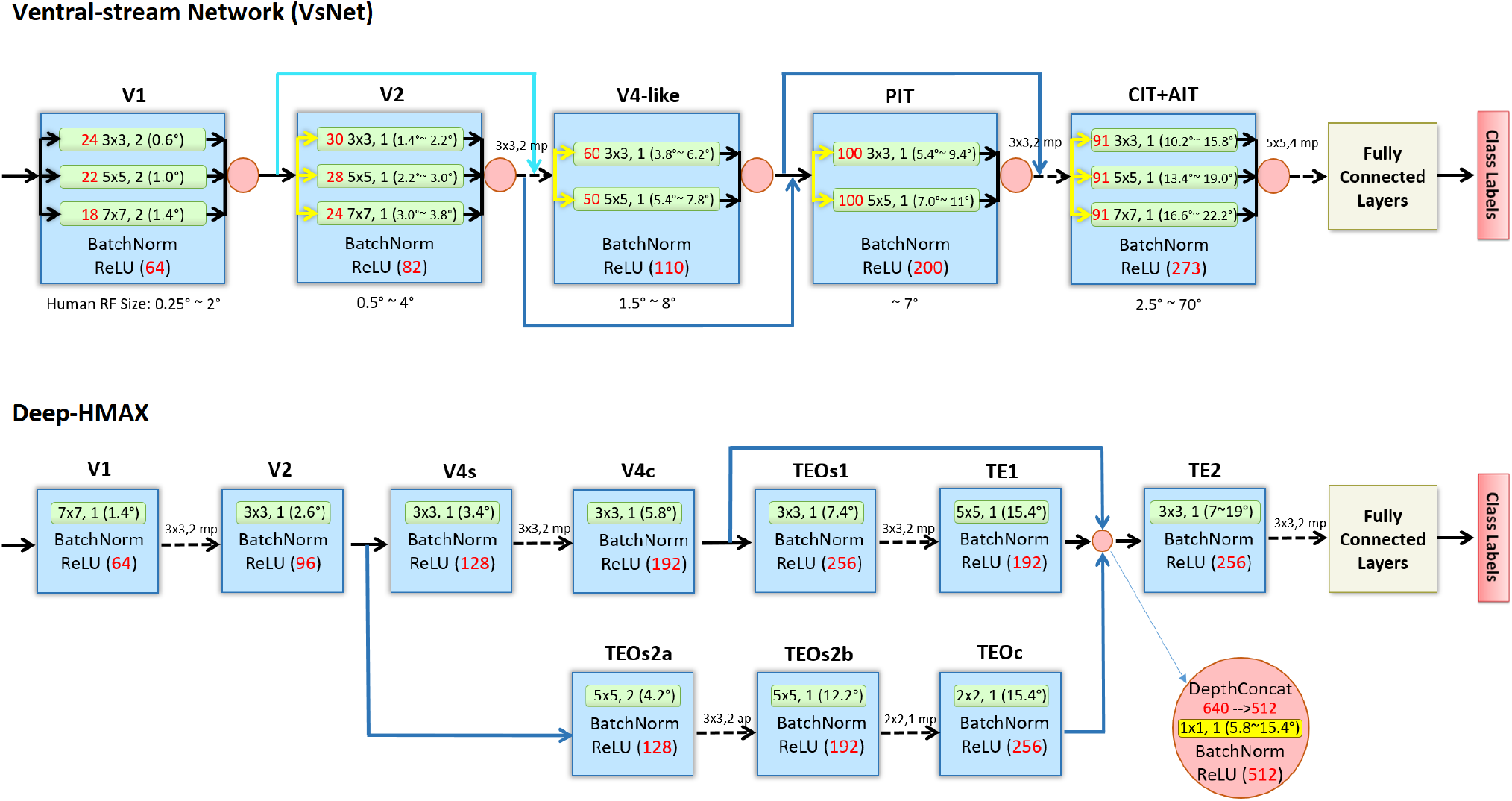
The architecture of our VsNet and Deep-HMAX designs. VsNet: each blue box represents a convolutional layer, with the corresponding ventral-pathway area labeled above. Pink circles are Depth-Concat layers that concatenate the input maps from the depth dimension. Arrows indicate input to output direction, dashed arrows represent max-pooling layers and their kernel sizes and strides, yellow arrows represent dimensionality reduction via 1×1 filters, and blue arrows are skip connections which is either a direct copy (dark blue), or a dimensionality reduced copy (light blue) via 1×1 filters. Green rectangle within each layer represent a set of filters, where the number of filters is in red, followed by the filter size and the stride size, with the corresponding receptive field (RF) size in visual angle shown in parentheses (assuming 1° spans 5 pixels). Note that the design of both VsNet and Deep-HMAX has the RF sizes of the convolutional filters in each layer to be as similar as the range of the RF size estimates in each of the five human ventral visual pathway areas. These target RF size ranges are indicated at the bottom of each VsNet layers (please refer to Supporting Information (SI) for the details of how these estimates are obtained). Each convolutional filter is followed by a Batch Normalization layer (BatchNorm) [21] and a Rectified Linear activation layer (ReLU). For more detailed architecture specification, please refer to SI.

Our CNN-CCF extraction algorithm is general, and can be applied to the filter responses from any pre-trained CNN. This makes model comparison possible. In addition to extracting CNN-CCFs from VsNet, we used the identical algorithm to extract CNN-CCFs from two other deep networks. One of these was AlexNet [30], a widely used CNN also consisting of five convolutional and two fullyconnected layers. Although AlexNet’s design was not brain-inspired, it has been used with good success in recent computational neuroscience studies [26, 4, 18] and is therefore of potential interest. More fundamentally, it will serve as a baseline against which the more brain-inspired networks can be compared, which is important to gauge broadly how the inclusion of neural constraints in a CNN’s design translates into improved prediction performance. We also extracted CNN-CCFs from a model that we are calling Deep-HMAX, a CNN version of the influential HMAX model of object recognition [53]. HMAX was designed to be a biologically plausible model of how the recognition of visually complex objects might be implemented in ventral brain circuitry [48, 60], but it was designed with handcrafted filters and therefore cannot be fairly compared to more recent and powerful convolutional network architectures. Our Deep-HMAX model keeps the basic architectural design elements of HMAX intact, central among these is the inclusion of simple and complex cell units, but replaces the originally hand-crafted units with convolutional layers that learn the simple and complex cell responses from visual input, thereby making possible more direct comparison to VsNet. Figure 2 shows the architecture of Deep-HMAX, and MM should be consulted for more details. Broadly, the model has a very different architecture than VsNet, with one example being its 10 convolutional and two fully-connected layers. This makes a comparison between VsNet and Deep-HMAX potentially valuable as a means of exploring the gross level of brain organization that should be designed into artificial deep neural networks. Critically, in our model comparison VsNet was computationally disadvantaged in that it used the least number of convolutional filters to predict attention control; AlexNet has 1152 filters, Deep-HMAX 1760, but VsNet only 726 (excluding 1×1 dimensionality-reduction filters). This conservative design means that, to the extent that VsNet better predicts attention control than the other models, this benefit is likely due to its brain-inspired architecture rather than sheer computational power.

## 3. Results

### CNN-CCF Predicts Visual Attention Control

VsNet, AlexNet, and Deep-HMAX were trained using ImageNet [50], then fine-tuned using the SBU-68E dataset (https://github.com/cxy7452/CNN-CCF/tree/master/SBE-68E/): an image dataset collected for this work consisting of 48 subordinate, 16 basic, and 4 superordinate categories, with each category having 550 image exemplars. The network was fine-tuned by using training/validation splits of 500 and 50 images, respectively, per category. (see MM for training details).

Features were extracted from the five convolutional layers of each identically-trained network using the previously described CNN-CCF feature selection method, and the number of CNN-CCFs were determined for each model. Recall that Yu et al. (2016) found that the number of BOW-CCFs extracted from their model accurately predicted the time that participants took to first fixate a target category cued at each of the three hierarchical levels. Our first evaluation was therefore to see whether CNN-CCFs from the three deep network models were as successful in predicting the hierarchical level of the target category. As shown in Figure 3A, all of the models tested were highly successful in capturing the trend of increasing time-to-target with movement up the category hierarchy in the behavioral data from Yu and colleagues (2016; see MM for additional details about the behavioral methods and data). This demonstration is important in showing that the number of CCFs is highly generalizable in its ability to predict the effect of hierarchical level on categorical guidance; more CCFs can be extracted for subordinate-level categories compared to basic, and for basic-level categories compared to superordinate, and these greater numbers of features form better target templates that can more efficiently guide attention to the categorical targets.

**Figure 3.**
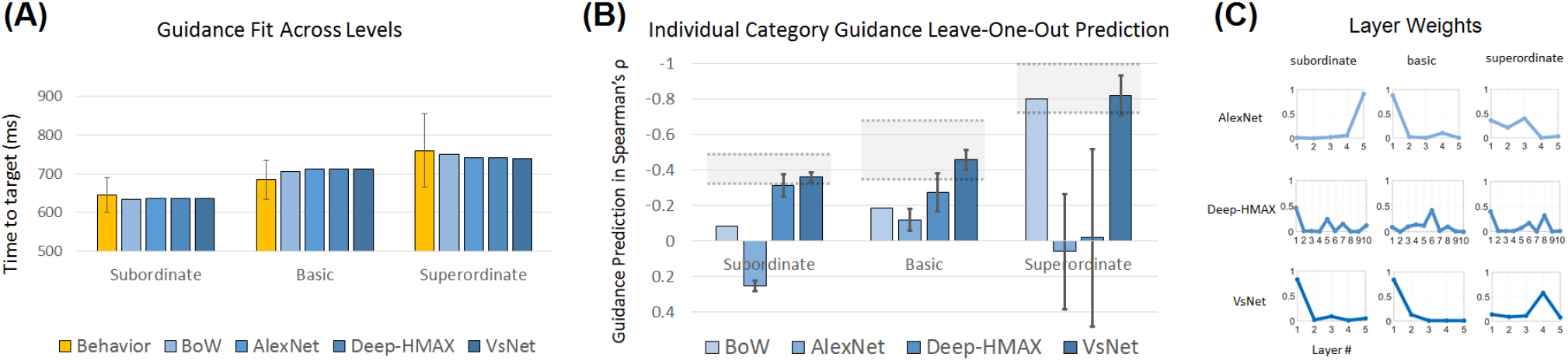
A. Human attention control (time to fixate the target) and the performance of one CCF model using Bag-of-Words (BoW) and three CNN-CCF models: AlexNet, Deep-HMAX, and VsNet. Results are grouped by hierarchical level, and the models performances are linearly scaled to best fit the behavior (thereby putting the models’ results in the behavioral scale). All four models are able to predict the subordinate-level advantage in attention guidance to a categorical target. B. Performance of the four models in predicting attention control to individual target categories within each hierarchical level, evaluated using the leave-one-out (LOO) method. Given the inverse correlation between number of CCFs and the time needed to guide attention to a target ([70]), more negative correlations indicate better predictions of attention control. Grey regions indicate performance ceilings in how well a model can predict attention guidance, defined by +/− one standard deviation of the mean guidance from a “subject model”. The subject model was also computed using LOO, only now we found the Spearman’s *ρ* between *n* − 1 participants and the participant who was left out (the mean and standard deviation was obtained by repeating this for all participants). The results show that a model using AlexNet to extract CCFs is unable to predict human behavior. However, the more brain-inspired CNN-CCF models are better, with VsNet being the best and on par with a human subject model. C. The best fitted weights by convolutional layer for each CNN-CCF model, grouped by hierarchical level. VsNet’s weight distribution suggests that categorical guidance at both subordinate and basic levels is driven by low-level features, while guidance to superordinate-level categories is driven by high-level features.

Capturing the behavioral guidance trend across category levels is one thing, using CCFs to predict guidance efficiency to individual categories is a different and far more challenging goal. Our experimental logic, however, is the same; the more CCFs that can be extracted for a target category, the better attention should be able to guide gaze to an unseen exemplar of that category. Across categories we therefore predict a negative correlation between the number of CCFs and the search time-to-target measure, with more CCFs leading to shorter target fixation times. But recall that each layer of a network is extracting its own CCFs, and it is unreasonable to believe that the attention control mechanism would disregard network depth and weigh all of these features equally. Hence, our approach was to find a CCF weighting across each network’s convolutional layers that optimizes a correlation (Spearman’s *ρ*) between the number of CCFs extracted at each layer and time-to-target, our behavioral measure of attention control, with each network model having its own optimized layer weights. The advantage of this formulation is that it allows Spearman’s *ρ* to be used directly as an objective function to optimize the layerwise weights, *W*, which we did using beam search with random steps [62] (see MM for details).

Figure 3B shows these category-specific predictions of guidance efficiency at each hierarchy for the four tested CCF models. Note that prediction success is indicated by higher negative correlations, plotted upward on the y-axis. Predictions from the BoW-CCF model were poor for subordinate and basic level categories and significantly worse than those from VsNet and Deep-HMAX. A very good prediction was obtained at the superordinate level, but given that there were only four categories at this level a high correlation might simply have resulted from chance. Interestingly, the number of CNN-CCFs extracted from the widely-used AlexNet model failed entirely in predicting attention guidance to individual target categories. This result drives home the fact that not all CCN models are equal; if the goal is to predict human behavior, CNNs should be modeled after the brain. Of the two evaluated brain-inspired CNNs, prediction success from Deep-HMAX was not reliably different from VsNet at the subordinate level (*p* = 0.059), significantly lower than VsNet at the basic level (*p* < 0.001), and non-existent at the superordinate level, while VsNet’s predictions remained very good. Indeed, for individual categories at all three hierarchical levels, VsNet’s predictions were well within the performance ceilings (gray regions) computed by having n-1 participants predict the behavior of the participant left out. This means that VsNet’s predictions were as good as can be expected given variability in the participant behavior, and it is the only model of the four tested for which this was consistently the case. Collectively, these results suggest that not all brain-inspired CNNs are created equal; a CNN designed after the ventral visual pathway is preferred over the architecture of Deep-HMAX.

CNNs have been criticized as being “black boxes”; they perform well but the reason for their success defies understanding. We prefer to think of CCNs as “transparent boxes” (a term borrowed from Aude Oliva, personal communication on May, 2017; see also [1]), ones that we can probe and peer into in attempts to decipher how they work. As one example, Figure 3C plots the optimized layer weights (*W*), grouped by level in the category hierarchy, for each of the three CNN models tested. VsNet shows very similar weight distributions for subordinate and basic-level targets, one clearly dominated by early features in its predictions of attention control. However, for superordinate category targets the CCFs learned by its CIT+AIT layer were the most predictive. Speculatively, this suggests that lower-level features may be driving attention control when relatively clear visual properties of the target can be inferred (e.g., object edges/rigidness/shapes and colors, depending on the category), but that higher-level features must be used for superordinate-level targets that do not have clearly representative visual properties. In contrast to this reasonable layer weighting, the optimized weightings for Deep-HMAX across its 10 layers seemed more erratic (although perhaps suggesting the emergence of a pattern similar to VsNet), and the optimized layer weights from AlexNet were uninterpretable since they produced low correlations. Once again, the brain-inspired design of the CNN appears to matter.

### CNN-CCF Visualization

Another example of VsNet being a transparent box is that it is possible to peer inside to see what patterns in images its CCFs were coding–the representative visual features of an object category. The CCFs for a given category can be visualized by finding the regions in input images that best activate a given CCF (a particular convolutional filter). Specifically, we first forward-pass an image of a category exemplar through VsNet to obtain the maximally-responsive locations in a feature map for the CCF of interest, and then probe backwards from the filters most activated location to the pixels in the image that was causing this maximal response [72] (see SI for the detailed algorithm). Figure 4A visualizes the image regions eliciting the five largest responses from CCFs, based on sorting their SNR scores, at each of VsNet’s layers for one exemplar image from the taxi category. Striking is the fact that these maximaly-active CCFs seem in some cases to be representing object parts that are specific to typical taxis, such as the signs attached to the roofs, but also parts that are more broadly representative of cars, such as wheels, windows, and side mirrors. This observation also nicely illustrates the generative nature of CCFs; they code the features that are common to a category (rooftop advertisements and wheels in the case of taxis) regardless of whether they are discriminative (police and race cars also have wheels, but only taxis have rootop signs).

**Figure 4.**
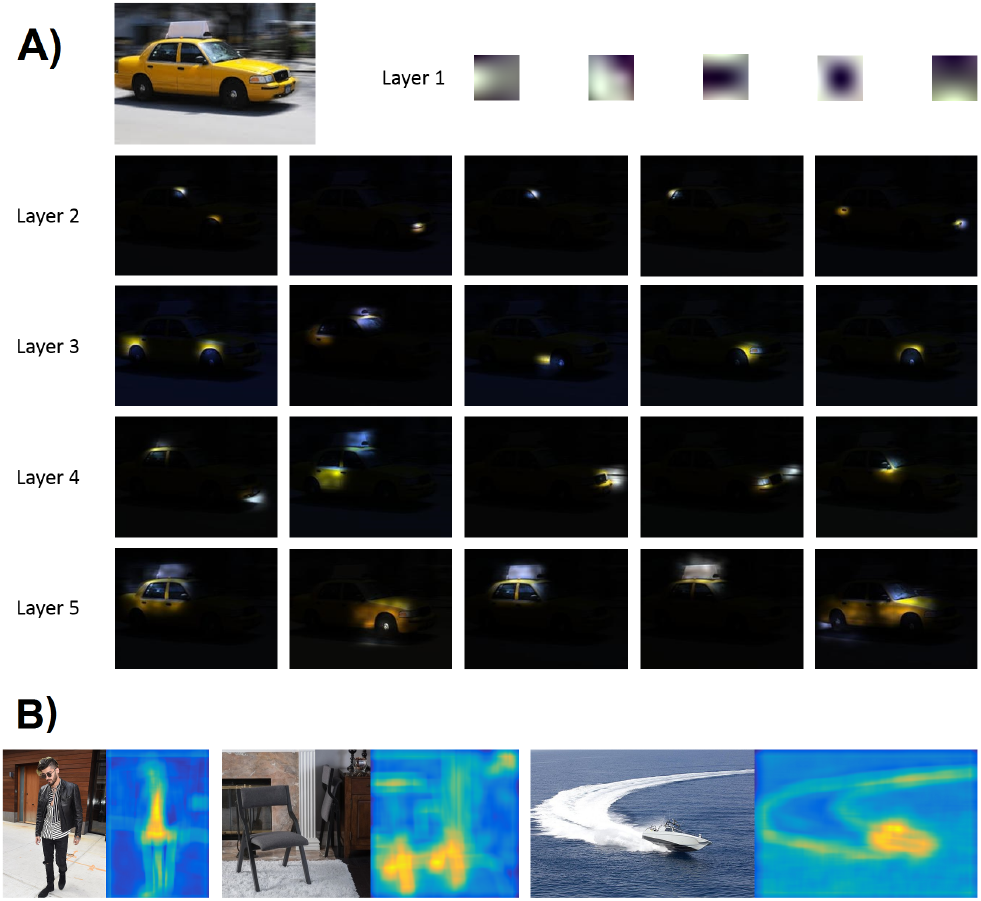
A. The CNN-CCF visualization of the taxi cab category. The visualized patches are the top 5 CCFs based on their SNR of each convolutional layer in VsNet on an example taxi image. The CCFs are showing some clear parts that are indicative of a typical taxi, which includes tires, headlights, windows, and the taxi sign. B. examples of CNN-CCF as object detectors. The heat maps are the combined activations of the given category’s CNN-CCFs, where the brighter the more activations. Categories from left to right: shirt (basic), folding chair (subordinate), and speedboat (subordinate).

The aggregated locations of maximally-active CNN-CCFs can also be used to detect categories of objects. This is because these CCFs will be broadly capturing the different parts of an object category, at different scales, making it possible to detect the presence of a target object in an image simply by detecting its constituent parts (CCFs). As qualitative examples, Figure 4B shows images depicting the object categories of shirt, folding chair, and speedboat, paired with the combined activation maps from CCFs extracted for those categories. None of these images were part of VsNet’s training set. Note that the CCFs for the shirt category precisely differentiate that object from the categorical siblings of jacket and pants, and that the CCFs for the category of folding chair are clearly coding the chair’s legs, which happens to be a part that discriminates that subordinate-level category from other chairs. The speedboat example is interesting in that it dramatically illustrates the difference between bottom-up saliency and top-down goal-directed attention control; CCFs activate strongly to the small boat but almost not at all to its far more salient white wake. What is significant about this demonstration is that this precise object localization was accomplished simply by combining the CCF activation maps without any additional processing costs.

### Large-Scale Image Classification

Prior to extracting CCFs from the three CNN models, the networks must be trained to learn an initial set of features. This initial training, and later validation, was done using ImageNet [50] following standard training procedures (see MM for additional details and subsequent fine-tuning). Although not directly within this study’s question of focus (goal-directed attention control), initial training results are an important indicator of how different network architectures grossly affect learned feature quality and network classification performance. Given their ready availability, we therefore discuss these results briefly here. Another motivation for raising this topic is that in our opinion VsNet produced interesting behavior, which we think might also be of interest to readers doing large-scale object classification.

Table 1 summarizes model performance on the validation set of the ImageNet dataset. Classification accuracies for all models were high, with validation errors indicating that the networks were successfully trained. However, VsNet achieved the lowest errors while AlexNet achieved the highest errors. This is an interesting finding because VsNet’s design was engineered after the ventral stream of visual brain areas, and was not designed or optimized for classification accuracy. Yet, it outperformed a model that was optimized for classification, AlexNet, by what is considered a significant margin in the computer vision literature. Moreover, although deeper CNN architectures generally outperform shallower networks [56], in VsNet we found an exception to this rule. While Deep-HMAX, the deepest network of the three, outperformed AlexNet in classification accuracy, it was less accurate than the shallower VsNet.

**Table 1.** Top-1 and top-5 validation accuracies for the three CNN models on the ImageNet dataset [50]. The overall high accuracies indicate the low likelihood of overfitting. Note that network performance improved with the degree of brain inspiration in its design.

| ImageNet | AlexNet | Deep-HMAX | VsNet |
| --- | --- | --- | --- |
| Top-1 Accuracy | 57.7% | 59.6% | **61.5%** |
| Top-5 Accuracy | 80.6% | 82.4% | **83.9%** |

Also notable is the fact that VsNet achieved the highest accuracies despite having fewer convolutional filters compared to Deep-HMAX and AlexNet (726, 1760, 1152 filters, respectively, excluding 1×1 dimensionality-reduction filters). Similar to network depth, the number of non-1×1 convolutional filters correlates highly with network performance in the computer vision literature (AlexNet [30] < VGG [56] < GoogLeNet [58] < ResNet [17]). VsNet is an example of a DNN outperforming networks having many more convolutional filters. This reversal of trend suggests that VsNet was able to learn better representations using fewer filters, with this greater convolutional kernel efficiency (Figure 5) pointing to a meaningful benefit of its brain-inspired design. Lesioning of VsNet to isolate the contributions of individual design decisions will be a direction of future work.

**Figure 5.**
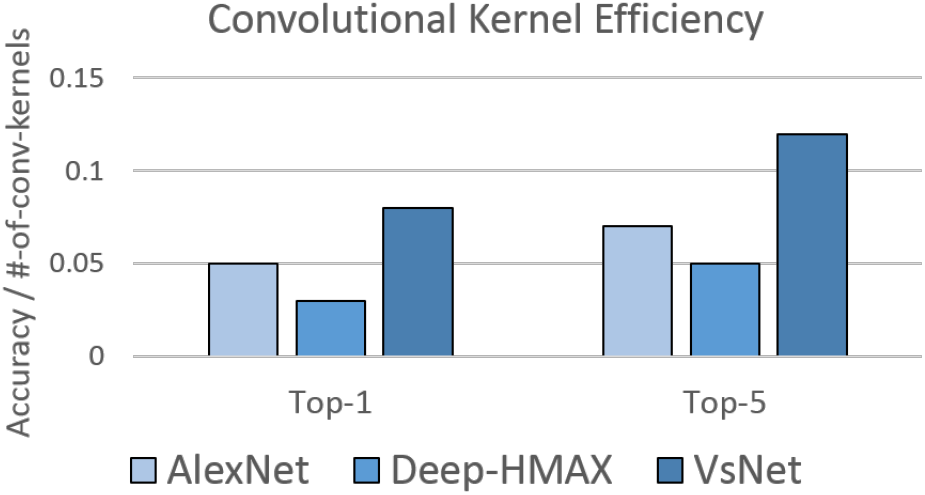
Plotted is a measure of the efficiency of a convolutional kernel, defined as accuracy per convolutional filter. VsNet was found to have the highest convolutional kernel efficiency, followed by AlexNet. Deep-HMAX was the least efficient network, possibly due to its long parallel branches learning redundant features.

## 4. Discussion

This study is the first to use a CNN to predict a goal-directed behavior, in our case the guidance of gaze during search. More specifically, we showed that an artificial deep neural network, one whose design is broadly informed by the architecture of the primate ventral visual stream, outperformed a less brain-inspired model (Deep-HMAX) and a popular yet brain-uninspired model (AlexNet) in predicting the control of attention to individual categories of common objects selected from a three-level category hierarchy.

Computationally, this demonstration is significant in two respects. First, VsNet outpredicted AlexNet, which had 58% more learnable filters, and Deep-HMAX which had 142% more. This is rarely observed, with the more common relationship being an increase in prediction success with the number of convolutional filters (AlexNet [30] < VGG [56] < GoogLeNet [58] < ResNet [17]). To our knowledge, this is the first time that a deep network with significantly fewer convolutional filters outperformed a more powerful network having many more. Second, given that deeper CNN architectures generally outperform shallower architectures [56], we were surprised that Deep-HMAX, a 10-layer CNN, did not compare more favorably to VsNet, a network half its depth. We speculate that this might be due to Deep-HMAX’s long parallel branches (forming after its V2 layer) creating a redundancy in the filters that are learned, in contrast to VsNet that uses short bypass connections to route lower-level information far more directly to higher-layers. Determining the specific sources of VsNet’s success, and the reasons why alternative brain-inspired designs fail, will be an important direction for future work. For now we can simply conclude that VsNet, a comparatively simple network in terms of its number of learnable filters, is learning representations of the target categories that yield superior predictions of human attention control in this search task. While a remarkable finding in one sense, in another it is not. If one assumes that the primate brain has already found a design and parameter specification to efficiently control the attentional routing of visual inputs through a (ventral) network of layers, it makes sense that a model’s predictive success might improve as its architecture becomes more like that of the brain.

The present work also extends our previous work on category-consistent features. We show that CCFs (generative and representative features of an object category) can be learned and extracted from CNNs, and that these more powerful CNN-CFFs successfully predict the degree that attention is guided to target categories. This is a potentially impactful finding, and in future work we will use CNN-CFFs to predict attention control in contexts other than visual search. Moreover, we showed that VsNet predicts guidance efficiency to individual target categories, and that its CNN-CCFs segregated reasonably across its layers, with lower-layers learning CCFs for subordinate and basic-level categories and higher-layers learning CCFs for more abstract categories. This generalization across categories demonstrates model robustness, and bodes well for the prospect of using the increasingly rich CNN-CCF feature representations to predict increasingly complex visual target categories.

Far from being a black box, VsNet hints at how search guidance and classification processes may interact across the layers of a deep network. A clue comes from the promising object localization made possible by projecting CNN-CCF activation from higher layers back to lower layers. Via this post-training backtracking method, the rich higher-level representations necessary for good classification reveal the spatial locations of lower-level target features coded by earlier visual areas, thereby delineating the object in space and creating the opportunity to bias that region for selective routing. Under this framework, guidance and classification are inextricably linked; better classification leads to stronger guidance, which in turn leads to better classification. We believe that understanding the role of object classification is essential to understanding attention control, as perhaps the greatest role of attention control is to make better classifications, mediated by an intelligent routing of visual inputs through the ventral stream.

## 5. Materials and Methods

### Behavioral Data Collection

Behavioral data were obtained from [70], and were collected using the SBU-68 dataset. This dataset consisted of crossly-cropped images of 68 object categories that were distributed across three levels of a category hierarchy. There were 48 subordinate-level categories that were grouped into 16 basic-level categories that were grouped into 4 superordinate-level categories. Stony Brook University undergraduates (n=26) participated in a categorical search task. On each trial a text cue designating the target category was displayed for 2,500 ms, followed by a 500 ms central fixation cross and then a six-item search display. Distractors were from random non-target categories and on target-present trials the target was selected from on of the 48 subordinate-level categories. Participants responded present or absent as quickly as possible while maintaining accuracy, and there were 144 target-present and 144 target-absent trials presented in random order. For each target-present trial, a participant’s goal-directed attentional guidance was measured as the time taken to first fixate the cued target. Please refer to [70] for full details of the behavioral stimuli and procedure.

### VsNet Design

VsNet is brain-inspired in three key respects: the number of filters at each convolutional layer is proportional to the estimated number of neurons in the corresponding brain structure, the sizes of filters at each layer are proportional to neuron receptive field sizes in corresponding structures, and the gross connectivity between its layers is informed by connectivity in the primate ventral visual stream. Each of these brain-inspired constraints will be discussed in more detail. With respect to VsNet’s broad mapping of convolutional layers to brain structures, it’s mappings between layer1 and V1 and layer2 and V2 are relatively noncontroversial. However, we wanted VsNet’s third convolutional layer to map to V4, a macaque brain area, and identifying a homolog to V4 in humans is less straightforward. A structure has been identified as “human V4” (hv4), and neurons in this structure are organized retinotopically [38, 11, 3] like macaque V4, but their feature selectivities are somewhat different. Macaque V4 neurons are selective to color, shape, and boundary conformation [7, 45, 5], whereas neurons in hV4 respond mainly to just color and occupy a proportionally much smaller cortical surface area [38, 3, 31]. For humans, shape and boundary and other object-related processing likely occurs in lateral occipital areas 1 and 2 (LO1/2) [31]. LO1/2 is also retinotopically organized and is anatomically adjacent to hV4 [10]. In an effort to obtain a sufficiently large number of learnable mid-level features, we therefore map VsNet’s third convolutional layer to a combination of hV4 and LO1/2, referred to here as “V4-like”. We intended VsNet’s deeper layers to map to IT, and decisions had to be made about these mappings as well. To keep congruence with the monkey neurophysiology literature, we specifically wanted to identify human homologs to macaque TEO and TE. For VsNet’s fourth layer we settled on a structure anterior to hV4, termed “human TEO” in [24, 25, 2] and PIT elsewhere [44], and for its fifth layer we chose central and anterior inferotemporal cortex (CIT+AIT) [46], roughly macaque TE.

### Ventral Stream Surface Areas

The numbers of convolutional filters in VsNet’s layers were based on estimates of human brain surface areas in the mapped structures. Specifically, V1, V2 and V4-like surface areas were estimated to be 2323 *mm*^2^, 2102 *mm*^2^, and 2322 *mm*^2^, respectively [31]. For PIT and CIT+AIT, we estimated their surface areas to be approximately 9 times larger than the surface areas in the corresponding macaque structures (TEO and TE, respectively) [44], based on reported differences in cortical size between macaque and human [10]. This resulted in an estimate of PIT having a surface area of 3510 *mm*^2^, and of CIT+AIT having a surface area of 3420 *mm*^2^. Having these surface area estimates, one approach might make proportional allocations of convolutional filters at each layer, but this would ignore the fact that some of these structure have a retinotopic organization. Retinotopy requires that the receptive fields (RFs) of neurons having similar selectivities are tiled across the visual field in order to obtain location-specific information, and this duplication of neurons is a major factor determining the surface area of some brain structures. CNNs have no retinotopy, their filters are convolved with a visual input rather than duplicated and tiled over an image. To equate the two, we derive a duplication factor that estimates the latent number of uniquely selective neurons within each brain structure, and then make the number of convolutional filters in the corresponding layer proportional to this estimate. In doing this we make a simple assumption. If the average RF size for a neuron type in a ventral stream structure is as large as the entire visual field, then there would be no need for the retinotopic duplication of this type of neuron for the purpose of capturing information from across the visual field. This would lead to a duplication factor of 1. However, if in this example the average RF size for neuron type covers only a quarter of the visual field, then there would minimally need to be four neurons of this type organized retinotopically to cover the entire visual field. This would lead to a duplication factor of 4. More generally, the following formulas were used to calculate the duplication factor for a given ventral stream structure and to determine the number of convolutional filters in VsNet’s corresponding layer:

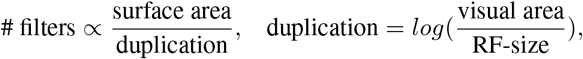

where both the area of the visual field and neuron RF size are expressed in degrees squared. We take the log of these values’ proportion in order to scale down the increase in the numbers of filters from lower to higher layers so as to stay within hardware constraints. For the current implementation, 1° of visual angle equaled 5 pixels, making the 224 × 224 pixel input images subtend approximately 45° × 45° of visual area. For each ventral stream area, we then take the average RF size at 5.5° eccentricity to be representative of neuron RF sizes in that structure (i.e., we currently do not capture here the foveal-to-peripheral increase in RF sizes, but see below). Doing these calculations, we obtained the representative RF size estimates of 1°, 3°, 5°, 7°, and 12° for V1, V2, V4-like, PIT, and CIT+AIT, respectively (see also [28]). Finally, using these values in the duplication factor calculation, and setting the total number of filters in the first convolutional layer (V1) to 64 (to be directly comparable to AlexNet), we obtain the final VsNet architecture consisting of 64, 82, 110, 198, and 272 filters for its 5 convolutional layers, excluding 1×1 dimensionality-reduction filters (see Figure 2).

### Receptive Field Size

In primates, the RFs of visually-responsive neurons increase in size with distance along the ventral stream; neurons in structures early in this pathway have small RFs, those in later structures have larger RFs [12, 28]. Moreover, within visual structures preserving retinotopy (V1 to V4) cortical magnification causes neurons coding the central visual field to have relatively small RFs, and neurons coding increasingly periphery locations to have increasingly larger RFs [9, 63, 12, 15]. VsNet was designed to grossly capture both of these properties. However, this latter relationship between RF size and visual eccentricity is difficult to implement in a CNN, where most models have filters of only a single size within each of their convolutional layers (i.e., [30, 72, 56, 17], with the exception of the Inception Module from [58]). This is because the convolutional filters in a CNN were specifically designed to not operate at specific image locations (shared weights), making the modeling of a changing retinotopy difficult. To approximate the variability in RF sizes due to scaling with eccentricity, we used parallel sets of 3×3, 5×5, and 7×7 pixel convolutional filters in each of VsNet’s layers (except for layers 3 and 4, which used only 3×3 and 5×5 filters). These sizes were chosen so as to approximate the range of RF sizes within corresponding structures, as estimated in [57, 25, 49, 15]. Assuming that 5 screen pixels correspond to 1 degree of visual angle, the 224×224 pixel ImageNet images used for training subtended a visual angle of 45°. More importantly, a 3×3 filter In VsNet’s V1 layer spanned 0.6°, a 5×5 filter spanned 1°, and a 7×7 filter spanned 1.4°. This range of RF sizes (0.6° to 1.4°) maps closely onto the range of RF sizes in V1 (0.25° to about 2°). These filters convolve with the input and then output feature maps that we concatenate in depth, such that the convolutional filters at the next higher layer (V2) receives responses from filters having three dfferent sizes. For example, stacking layer 2’s 3×3 filters on top of layer 1’s 3×3, 5×5, and 7×7 filters, results in layer 2’s 3×3 filters having RF sizes of 1.4°, 1.8°, and 2.2°, respectively (the parenthetical values listed in Figure 2 for VsNet’s 3×3 V2 filters). Doing this for the 5×5 and 7×7 filters produced a range of sizes again corresponding well to the range of RF sizes observed in V2 neurons (Figure 2). A similar procedure was followed for VsNet’s V4-like layer, which produced similarly good estimates of neuron RF sizes. Over VsNet’s first three layers, the filters at each higher layer therefore had, not only larger RFs, but also a broader range of RF sizes. For VsNet’s PIT and CIT+AIT layers, the same numbers of filters were allocated in the parallel sets, reflecting the relaxation of a retinotopic organization in the corresponding ventral structures. Notice that GoogLeNet’s Inception Module [58] also has this similar parallel-filter architecture, but it was not designed to be consistent to the primate visual cortex.

### Bypass Connections

In addition to the feed-forward projections that connect each ventral stream area with the next higher level along the pathway, good evidence also exists for connections that skip or bypass neighboring ventral structures [41, 23, 28]. VsNet captures both types of ventral stream connectivity, although it considered only a first-pass attempt to do so; capturing the minutia of this brain connectivity is currently beyond its scope. The direct connections are already embedded in its feed-forward design, so the focus here will be on detailing its bypass connections. Major bypass connections exist from V2 to TEO [41, 59] and from V4 to TE [59], with a weaker bypass connection known to exist between V1’s foveal region to V4 [41, 14, 61]. These three bypass connections were designed into VsNet. We added a weak bypass connection from layer 1 (V1) to layer 3 (V4-like), a full bypass from layer 2 (V2) to layer 4 (PIT), and another full bypass from layer 3 (V4-like) to layer 5 (CIT+AIT). We implemented these bypass connections by concatenating in the depth dimension of the output from the lower layer to the target layer’s input. Note that this concatenation method is different from the summation method used by ResNet [17], but is conceptually similar to the Inception Module design used by GoogLeNet [58]. Following [58], we also use 1×1 filters before each of VsNet’s convolutional layers (except layer 1, where they are not needed) for dimensionality reduction and memory conservation (yellow arrows in Figure 2). We chose this concatenation method in order to give VsNet maximum flexibility in how bypassed information is best combined with information at the target layer, which we believe is preferable to assuming that the cortex simply sums this information. Specifically, a full bypass was implemented by concatenating in the depth dimension a complete copy of the source layer’s output feature map to the end of the target layer’s input map. We implemented a weak bypass similarly, but now the source layer’s output map was depth-reduced (dimensionality reduced by half via 1 × 1 convolutional filters) before being concatenated with the target layer’s input feature map. Please refer to Table XXX for the detailed description of VsNet’s architecture.

### ImageNet Training

VsNet, AlexNet, and Deep-HMAX were trained using ImageNet. All training and validation images were resized to have the shortest side be 256 pixels while keeping the original aspect ratio, and the standard data augmentation methods of random crops (224×224) and random horizontal flips were employed. Center crops were used to compute validation accuracies at the end of each training epoch. The training batch-size for AlexNet, Deep-HMAX, and VsNet was 128, 64, and 60, respectively. Each network was trained using 4-threads with image data stored on a solid-state drive (SSD), and 60 training epochs took roughly 2 to 4 days to complete using a 2.93 Ghz Intel Xeon x3470 processor with 32 Gb of memory and a single Titan X GPU. Networks were implemented using Torch7, and the method from [16] was used for parameter initializations. Following ImageNet training, networks were fine tuned using the SBU-68E dataset, an expanded version of the SBU-68 dataset. The original SBU-68 dataset contained contained 4,800 images of objects, which were grouped into 100 exemplars from each of 48 subordinate-level categories [70]. These images were further combined hierarchically to create an additional 16 basic-level categories and 4 superordinate-level categories, yielding 68 categories in total. The expanded SBU-68E dataset built on the earlier dataset by exploiting Google, Yahoo, and Bing image searches to obtain 1,500 exemplars from each of the same 48 subordinate-level categories, thereby making it suitable for deep network training. GIST descriptors [42] were used to meticulously remove image duplicates, followed by a manual pruning of the images to ensure that those with incorrect class labels were removed and that the retained images were well-cropped around the labeled object. These exclusion criteria yielded 500 training and 50 validation images per category, for a total of 24,000 training and 2,400 validation images in the expanded set. All images were resized such that the shortest side was 256 pixels wide while retaining the original aspect ratio. See SI for more details on multi-task fine-tuning of the networks.

## Acknowledgements

Invaluable feedback was provided by the Computer Vision Lab and the Eye Cog Lab at Stony Brook University, and by Dr. Talia Konkle and the members of the Harvard Vision Sciences Lab. This work was also supported in part by a GPU donation from NVIDIA

## Supporting Information

### Randomized beam search with Spearman’s *ρ*

A basic problem is encountered when comparing CNN-CCFs extracted from a deep network to behavioral responses; because there are a different number of filters in each convolutional layer, how do we weight the CNN-CCFs extracted across these different layers when predicting behavior? Using VsNet to better illustrate the problem, it extracts a number of CNN-CCFs at each of its five convolutional layers (e.g., 5, 10, 12, 3, 7) for each of the 48 subordinate target object categories (e.g., “police car”). However, our behavioral measure of attention control is median time-to-target, a single value observed from each participant for each target category. To correlate the two we must therefore convert a predicted vector of size 48 × 5 (number of CNN-CCFs extracted per layer for each of 48 categories at the subordinate level) to a 48 × 1 vector of median behavioral guidance times (one measure for each of 48 categories). One method of doing this would be to simply sum CNN-CCFs across the five network layers, but this would make the questionable assumption that the brain treats the contribution of all layers equally when controlling attention guidance. Another method would be to use a linear model of the form *y* = Σ *w_i_x_i_*, where *y* ∈ ℝ and each *w_i_* denotes a weight attached to the number of CNN-CCFs extracted at the *i*^th^ network layer. Such linear models are usually optimized using least squares to best fit a target vector, but the traditional least-squares method cannot be optimized to minimize correlation loss, as Spearman’s *ρ* is the metric that we use to evaluate the CNN-CCF prediction quality to human guidance performances. Broadly, our goal is to find a set of weights that produces the highest Spearman’s *ρ* between the predicted vector, **Y**, and our measure of attention control, **Y’**. Specifically, we want to find a set of non-negative weights across a network’s layers such that the guidance prediction *y_i_* of the *i*^th^ category is of the form:

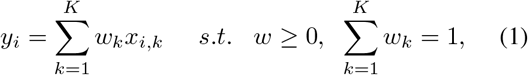

where *x_i,k_* is the number of CCFs extracted from the *k*^th^ layer of category *i*, and *w_k_* ∈ **W** is the weighted contribution of the *k*^th^ layer, where *K* = 5 for AlexNet and VsNet, and *K* = 10 for Deep-HMAX. We then want to evaluate the model’s output, prediction vector **Y**, and the target vector, median time-to-target **Y’**, so as to find a correlation between the inverse number of CCFs (1/**Y**) and guidance efficiency that optimizes Spearman’s *ρ*.

To find this optimized Spearman’s *ρ*, and therefore the best **W** for each network, we use the Randomized Beam Search (RBS) method from [62]. The procedure is as follows. We first randomly initialize *w_i_*S to non-negative values that sum to one, then compute an initial goodness-of-fit using this **W** and Spearman’s *ρ*. Following this initialization, RBS iterates through four operations: (1) an *i* is randomly selected and a new non-negative value for *w_i_* randomly generated, followed by re-normalization of **W** so that **W** sums to one, (2) a new goodness-of-fit is computed using the new **W**, (3) if this new **W** results in a better fit (higher Spearman’s *ρ*), keep the new **W**, else, revert back to the previous **W**, and (4) repeat steps 1–3. In the current implementation, we performed 300 of these iterations for each fit, and performed 200 fits to find the best **W**. In piloting we found that this method converges usually within 200 iterations. We are therefore confident that our use of 300 iterations with 200 repetitions resulted in a highly optimized **W**.

Our goal is not just to fit data, but rather to determine how well an optimized weighting of CNN-CCF number across the convolutional layers of each network can *predict* attention control to a category goal. To do this, we obtained for each network its predicted **Y** over the n categories using a series of leave-one-out jacknife operations. For a given network, its **W** was fitted using *n* − 1 categories in order to obtain its highest correlation with participants’ median time-to-target, and then this optimized **W** was used to predict a *y* for the held-out category as specified in Equation 1. We repeated this procedure *n* times to obtain a complete prediction vector of **Y** over all tested categories, and then repeated this ten times to obtain a variance in order to evaluate the robustness of each model’s predictions.

### CNN-CCF Feature Visualization

In order to visualize the regions in an image that a CNN-CFF feature (a convolutional filter) best responds to, a correspondence between that filter and those pixels in an image must be established. Here we detail the non-trivial sequence of steps needed to make this correspondence.

The process begins by forwarding an image into a network and obtaining the feature map for the given convolutional filter that we wish to visualize. The largest 95% of values on this feature map are retained and the rest are set to zero. Broadly, the remaining steps involve working backwards (*backtracking*) through the series of convolutions that produced this pattern of feature map activation at a given network layer, to the specific pixels in the input image. To do this, we needed to perform inverse operations for several computations (convolution, max pooling, ReLU, etc.), not all of which are perfectly reversible. The resulting visualizations are therefore only approximations of the filter-specific network activations. Nevertheless, we see this as a useful tool for probing the “black box” and generating hypotheses to test in new experiments.

The specific steps in backtracking are as follows. (1) We perform *flipped convolution* [72, 73], meaning we flip the filters vertically and horizontally and convolve the feature map to get its input. (2) *Inverse max pooling*. During the feed-forward processing we record the the locations of the values that are taken during max pooling (the “switches”) [72, 73], making possible the inverse operation of restoring the values to their original positions and setting the other values to zero. (3) *ReLU*. The inverse of a ReLU operation is also a ReLU operation [72, 73]. (4) *Inverse batch normalization*. Having recorded the mean and standard deviation of the feature maps processed by each filter, the inverse operation of batch normalization involves multiplying each filter’s feature map by the map’s standard deviation and then adding the mean. (5) *Inverse average pooling with stride s*. This operation involves upsampling the image by a factor of *s*, and then filling the upsampled image locations with nearest values from the downsampled image. (6) *Inverse concatenation (detachment)*. Concatenation is used at multiple places in VsNet’s design, with one of these being in its bypass connections. For example, in the V2 bypass connection, the output of V1 layer is concatenated with the input to the V4-like layer. Reversing this operation requires detaching the data volume into 2 parts, one backtracked through the bypass over V2 and the other backtracked through V2. (7) *Inverse branching*. Using another example, the input to V2 is convolved by different numbers of filters of different sizes (3×3, 5×5 and 7×7 pixels) into 3 separate branches; to invert this operation these outputs must be summed over the 3 branches using flipped convolution [72, 73].

Using these inverse operations, we backtrack from the feature map generated by a target CNN-CCF filter to obtain the corresponding network input, what we refer to as the *projected image*. We then take the absolute values in the projected image at the input layer, sum these over the RGB channels, and normalize the summed values into [0,1] to obtain a heat map. Mapping the heat map onto the original image highlights the region in the original image that best activated the target kernel (Figure 4B).

### Fine Tuning the Networks

The SBU-68E dataset is hierarchical, and as such has multiple class labels for each category exemplar (i.e. a “passenger airplane” is also an “airplane” at the basic level and a “vehicle” at the superordinate level). We therefore fine-tuned the networks using a multi-task learning regime. We first replaced the single classification layer with three branches of newly initialized fully-connected classification layers, each corresponding to the 48 subordinate, 16 basic, and 4 superordinate output classes used in [70]. In doing this we used the same softmax layer and negative log-likelihood cross-entropy loss methods, but the backpropogation of loss down each of the three branches was given weights of 0.7, 0.24, and 0.06, respectively. These weights correspond to the proportion of object categories at each classification branch. Fine-tuning was performed using the same training settings as was used for the ImageNet pretraining, except for there being just 25 epochs and a reduced learning rate. Also different from pre-training, fine-tuning took just 3 hours to finish for all three networks, with all networks converging extremely well. Analyses revealed that top-1 errors were low, and that differences in errors across the models were small and likely not meaningful (less than 10% top-1 error for all networks at all three hierarchies). This is evidence that the fine-tuning was successful, and that the models generalized nearly perfectly to SBU-68E’s unseen validation set without any apparent over-fitting.

